# Explore the Potential of a Plant Phospholipase as an Antimicrobial

**DOI:** 10.1101/2020.10.17.343541

**Authors:** Curtis Chen, Shanhui Xu, Yanran Li

## Abstract

Global public health is increasingly threatened by the fast emergence of antibiotic resistance, and novel types of antibiotics are urgently needed. Metazoans have evolved their own antimicrobial mechanism, such as human group IIA secreted phospholipase A (sPLA2), which can efficiently inhibit the growth of gram-positive bacteria, but with much lower efficiency toward gram-negative bacteria. Here, we verified the antibacterial activity of a plant lipase, PLIP1 from *Arabidopsis thaliana*, against the gram-negative bacteria *Escherichia coli*, which belongs to the WHO priority 1 (critical) pathogen *Enterobacteriaceae* family. We also explored the potential of evolving PLIP1 as a more potent antimicrobial agent towards *E. coli*. Our results imply the possibility of using plant lipases as a potential antimicrobial and shed light on the future exploration of plant enzymes for novel and more efficient antibacterial agents.

## Introduction

Antibiotics are essential to public health, providing cures to bacterial infections [5]. However, due to the rapid emergence of antimicrobial resistance and the lack of new types of antibiotics, viable options to treat bacterial infections are decreasing, especially those to treat gram-negative bacteria [5]. The World Health Organization (WHO) recently highlighted the threat of gram-negative bacteria like *Acinetobacter baumannii, Pseudomonas aeruginosa*, and *Enterobacteriaceae*, which are often resistant to many antibiotics and can quickly evolve or acquire resistance [10]. Alternative mechanisms to treat these bacteria are desired.

Metazoans have evolved multiple strategies to combat microbial infections. One strategy is to use antimicrobial peptides and enzymes to disrupt the microbe envelop, e.g. human tears contain a number of antimicrobial agents such as lysozymes and group IIA secreted phospholipase A2 (sPLA2) [3]. sPLA2 can be purified from *Escherichia coli* [8], and has been identified as the major antimicrobial tear component against Gram-positive bacteria [6], yet exhibits low activity towards gram-negative bacteria. The antibacterial activity of sPLA2 is attributed to the fact that it can efficiently hydrolyze phosphatidylglycerol (PG) [6], which is the major lipid component of most Gram-positive bacteria. In gram-negative bacteria *P. aeruginosa* and *Enterobacteriaceae*, Phosphatidylethanolamine (PE) is more abundant [2]. Thus, a lipase that can efficiently degrade PE may serve as an antimicrobial agent against these deadly bacteria for external use, similar to sPLA2 in tears. Plants encode a large number of lipases. Taking the model organism *Arabidopsis thaliana* as an example, there are over 300 genes that are annotated as lipases [4], with more than 38 annotated or characterized as phospholipases. Thus, plants serve as a great resource for alternative lipases that exhibit higher hydrolysis efficiency towards various phospholipids. One recent study on a plastid lipase1 (PLIP1) from *A. thaliana* reported that *E. coli* is barely viable when PLIP1 was expressed, and that PLIP1 is efficient in catalyzing the hydrolysis of PE [9]. Here, we used PLIP1 to explore the antimicrobial activity of plant lipases and the potential of using plant lipases as antimicrobials for external utilization. In order to bypass the detrimental effect that PLIP1 has on *E. coli*, we expressed PLIP1 in yeast, which is not affected by this enzyme. Furthermore, since yeast is a eukaryotic cell, it may also simulate the effect that the PLIP1 enzyme will have on human cells. We were able to detect the inhibition of PLIP1 on *E. coli*, and tried to use the recently developed continuous evolution method OrthoRep [7] and adaptive evolution to engineer PLIP1 but without obtaining mutants of obviously enhanced activity, possibly due to the relatively low activity of PLIP1. Our results confirm the antimicrobial activity of PLIP1 towards gram negative bacteria when applied exogenously. However, extensive engineering work still needs to be performed to enhance the antimicrobial activity of PLIP1 so it can be utilized as a potent antimicrobial agent.

## Results and Discussion

### PLIP1 exhibits clear inhibition of *E. coli* growth

PLIP1 is originated from the chloroplast in *Arabidopsis thaliana* for the degradation of phospholipids [9]. PLIP1, classified as phospholipase A_1_, cleaves acyl groups from the glyceryl moiety at the sn-1 position, while human sPLA2 is a A_2_ phospholipase cleaving at the sn-2 position. Similar to sPLA2, PLIP1 is very likely to function as antimicrobial protein for external usage. Two well-documented secretion peptides, prepro-α and preOST1 [1], were used to secret the lipases from yeast cell into the medium by fusing the signal peptides to the N-terminus of the lipases. Low-copy number plasmids were used for the heterologous expression of the two lipases under the control of the strong *GPD* promoter. We also noted a clear growth deficiency of the yeast strain when PLIP1 was expressed from a high copy number plasmid, which indicates that in the presence of PLIP1 at higher concentration, yeast membrane will also be affected.

To examine the antimicrobial activity of PLIP1 towards *E. coli*, 2 μL of the PLIP1-serecting yeast culture (OD_600_ around 14.0) was dotted onto petri-dishes with a “lawn” of *E. coli* on top and allowed to grow for 48 hours. Compared to the negative controls (wild-type yeast strain), visible inhibition zones were observed surrounding the PLIP1-secreting yeast strains (Figure 1A). To further confirm the inhibition activity of PLIP1 towards *E. coli*, we cultured *E. coli* in the 48-hour medium of yeast secreting either PLIP1 or the inactive PLIP1 mutant (PLIP^S422A^). By measuring the cell density (OD_600_) of *E. coli* over a time course of 24 hours, we found that the medium of the PLIP1-screting yeast clearly inhibited the growth of *E. coli*, in comparison to PLIP^S422A^-producing yeast strain (Figure 1B). Surprisingly, we do not see the inhibition activity of preOST1-PLIP1, which is likely due to the less efficient secretion efficiency of preOST1 in the case of PLIP1. The prepro-α-PLIP1-secreting yeast was therefore used in the subsequent investigations.

**Figure 1.**
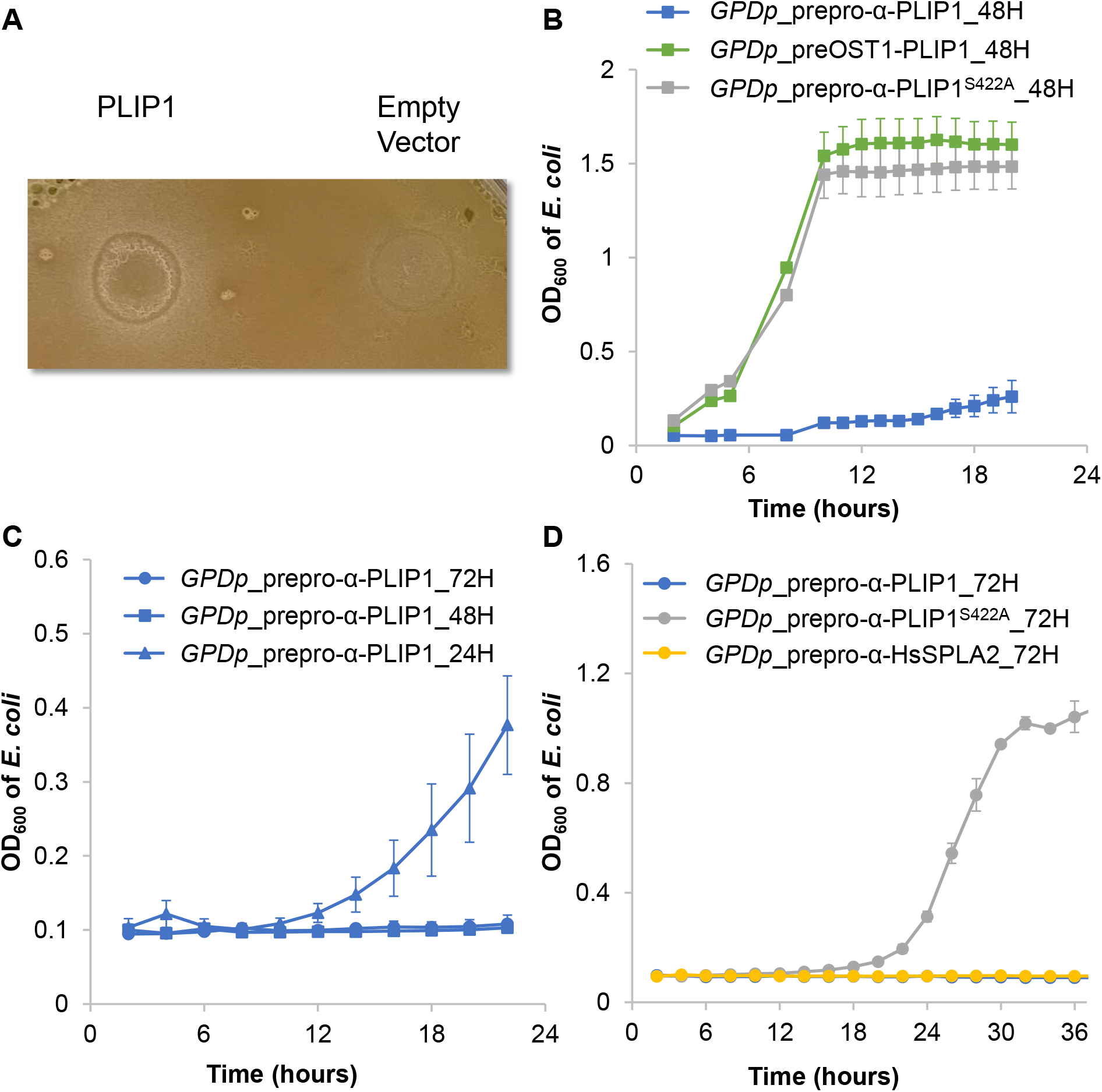
Characterization of the inhibition activity of PLIP1 on *E. coli*. **(A)** Agar diffusion assay of yeast strains harboring (1) a low-copy number plasmid encoding prepro-α-PLIP1 downstream of *GPD* promoter, (2) empty vector on a lawn of *E. coli*. The plate was incubated at 30 °C for two days. **(B)** Growth curves of *E. coli* in the 48-hour spent medium of yeast strains expressing prepro-α-PLIP1^S422A^ (grey), pre-OST1-PLIP1 (green), and prepro-α-PLIP1 (blue) on a low-copy plasmid under the regulation of *GPD* promoter. **(C)** Growth curves of *E. coli* in the 24 (triangle)-, 48 (square)-, and 72 (circle)-hour spent medium of yeast strains expressing prepro-α-PLIP1 on a low-copy plasmid under the regulation of *GPD* promoter. **(D)** Growth curves of *E. coli* in the 72-hour spent medium of yeast strains expressing prepro-α-PLIP1^S422A^ (grey), pre-OST1-PLIP1 (green), prepro-α-PLIP1 (blue), and prepro-α-HsSPLA2 (yellow) on a low-copy plasmid under the regulation of *GPD* promoter. All data points are representative of at least three biological replicates for each *E. coli* culture and the error bars represent the standard deviation of the replicates.

To check whether 48-hour culturing is enough for the PLIP1-secreting yeast strain to produce enough PLIP1 into the medium, we examined the growth of *E. coli* in the yeast medium cultured for 24, 48, and 72 hours after back dilutions. Our results indicate that the 24-hour medium showed no inhibition on *E. coli*, while the 48- and 72-hour medium showed clear inhibition with 72-hour medium slightly more efficient in inhibiting *E. coli* (Figure 1C). Furthermore, to compare the inhibition activities of PLIP1 and the human lipase SPLA2, the antibacterial effects of which has been well documented, *E. coli* was cultured in the 72-hour medium of PLIP1 and sPLA2-screting yeast strains. The cell density (OD_600_) of *E. coli* was measured for 24 hours, using the inactive PLIP1^S422A^ mutant as a negative control. Our results indicate that the inhibition activities of PLIP1 and the human lipase SPLA2 are comparable (Figure 1D).

### Mutation of PLIP exhibits higher potency towards E. coli

The OrthoRep system is a recently developed continuous evolution method harnessing an orthogonal DNA replication system that can stably mutate the target gene 10,000-fold faster than the natural mutation rate of the genome [7]. The OrthoRep system thus allows us to mutate the PLIP1 gene in a targeted fashion with higher degradation activity towards *E. coli* cell membrane. We hypothesized that when *E. coli* and PLIP1-secreting yeast are co-cultured, the shortage of the nutrients will provide a driven force for the evolution of PLIP1 towards being more efficient in inhibiting the growth of *E. coli*. Unfortunately, the expression level of the gene of interest in the OrthoRep system has been optimized to ~ 40% of the strength of the *GPD* promoter [11], which is not enough for us to observe a clear delay in the growth of *E. coli* when we cultured *E. coli* in the 72-hour medium of yeast expressing PLIP1 using the OrthoRep system (Figure 2A).

**Figure 2.**
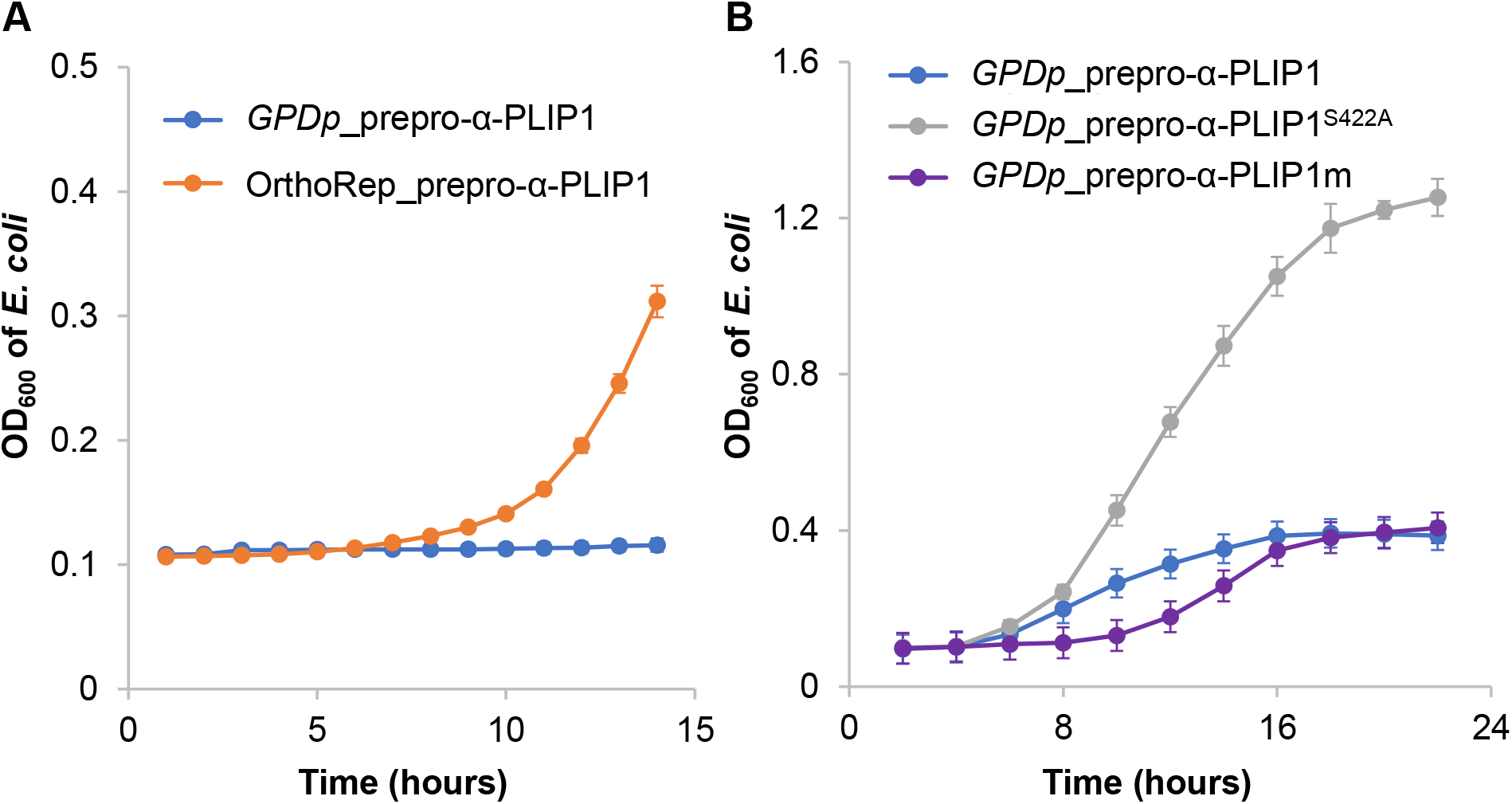
Optimization of PLIP1. **(A)** Growth curve of *E. coli* in the 72-hour spent medium of yeast strains expressing prepro-α-PLIP1 using the OrthoRep system (orange) and low-copy plasmid under the regulation of *GPD* promoter (blue). **(B)** Growth curves of *E. coli* in the 72-hour spent medium of yeast strains expressing prepro-α-PLIP1^S422A^(grey), prepro-α-PLIP1(blue)-, and prepro-α-PLIP1 mutant (purple) on a low-copy plasmid under the regulation of *GPD* promoter. All data points are representative of at least three biological replicates for each *E. coli* culture and the error bars represent the standard deviation of the replicates.

We then tried normal adaptive evolution instead. Over the course of two weeks, PLIP1-secreting yeast was co-cultured with *E. coli* for seventh rounds, with each round about 48 hours. At the end of each round, serial dilutions were repeated with the addition of fresh medium and fresh *E. coli* culture. The mutant from the 7th round showed slightly enhanced inhibition activity on the growth of *E. coli* (Figure 2B). The failure of obtaining a mutant with a substantial enhancement in the antibacterial activity towards *E. coli* is likely due to either the short timescales or the slower evolution.

In sum, we have confirmed the antimicrobial activity of PLIP1 towards gram-negative *E. coli*, showed the inhibitory activity of PLIP1 and the well-documented sPLA2 towards *E. coli* are comparable. We were not successful in obtaining a PLIP1 mutant of significantly enhanced antimicrobial activity. We believe that an OrthoRep system of the expression level further enhanced to a similar level to the strength of the *GPD* promoter will enable us to obtain such a mutant.

## Materials and Methods

### Materials and culture conditions

A haploid CENPK2.1D (*MATα; his3D; 1 leu2-3_112; ura3-52; trp1-289; MAL2-8c; SUC2*) was used in this work. Yeast strains were cultured at 30 °C in complex yeast extract peptone dextrose (YPD, all components from BD Diagnostics) medium or synthetic defined medium (SDM) containing yeast nitrogen base (YNB) (BD Diagnostics), ammonium sulfate (Fisher Scientific), 2% (w/v) glucose unless specified and the appropriate dropout (Takara Bio) solution for selection. *E. coli strain TOP10* was used in the inhibition assay, and cultured at 37 °C in lysogeny broth (LB) medium (Fisher Scientific).

### Plasmid and yeast strain construction

The PLIP1 gene was obtained from *Arabidopsis. thaliana* cDNA library. This sequence was then cloned using Polymerase Chain Reaction with Phusion high-fidelity DNA polymerase and annealed to the prepro-α or the pre-OST1 secretion signal peptide using Gibson Assembly. Gateway LR reactions were then used to assemble the secretion signal peptide-PLIP1 construct into the pAG414 plasmid vector. The recombinant plasmid DNA created from the gateway LR reactions were then transformed into Top10 competent *E. coli* cells. The *E. coli* cells were plated onto LB plates, and allowed to grow for 24 hours at 37°C. The presence of the plasmid was confirmed by performing colony PCR, using Genescript Taq Polymerase. Plasmids used in this study are listed in Table 1.

**Table 1.** Plasmids used in this study.

| Plasmid | Description |
| --- | --- |
| pYL134 | pAG414GPD-Prepro-α-PLIP1 |
| pYL153 | pAG414GPD-PreOST1-PLIP1 |
| pYL578 | pFDP-10PB-PLIP1 |
| pYL619 | pAG414GPD-Prepro-α-sPLA2 |
| pYL620 | pAG414GPD-Prepro-α-PLIP1-S422A |
| pYL621 | pAG424GPD-Prepro-α-PLIP1 |

### Inhibition assay of *E. coli* on solid agar medium

Yeast transformants or wild-type yeast strains were grown on SDM agar plates for 48 hours at 30°C. After this, single yeast colonies were picked and grown in liquid SDM medium for 48 hours. Simultaneously, wildtype *E. coli* was grown in liquid LB at 37°C overnight till stationary phase. 10 μL of the overnight *E. coli* culture was added to 100 μL sterile water and spread onto an agar plate (40 g/L of LB, 1 × SDM), and 2μL of the corresponding 48-hour yeast culture was applied onto the plate once the agar plate was allowed to dry. The plate was then incubated for 48 hours at 37°C. After 48 hours, inhibition zones were observed around the yeast colonies expressing PLIP1.

### Inhibition assay of *E. coli* using yeast cell culture supernatant

Yeast transformants were grown on SDM agar plates for 72 hours at 30°C. Then single yeast colonies were picked and grown in liquid SDM medium for 24, 48, or 72 hours. Simultaneously, *E. coli* was grown in liquid LB medium for 24 hours at 37°C. After the corresponding incubation hours, the yeast cells were separated from the medium using centrifugation at 13000 rpm for 10 minutes. 750 μL yeast cell-free supernatant, 200 μL 5 × LB broth, 50 μL MES buffer (pH 5.8), and 5 μL *E. coli* overnight culture (OD_600_ ~ 1.5) were mixed together in a new culture tube. The mixture was incubated at 37°C, and the cell density was monitored at OD_600_ until the growth curves of the *E. coli* had plateaued.

### Adaptive evolution of PLIP1 in yeast

*E. coli* and yeast containing the secretion signal-PLIP1 plasmid were grown together in 2 mL of combined SDM and LB liquid medium. 100 μL MES buffer (pH 5.8) was added to keep the pH constant. The culture was incubated at 37°C. After three days, the combined culture was diluted by extracting 20 μL and adding it to 2 mL of fresh mixed liquid medium. A SDM plate was restreaked using the remaining solution to ensure the survival of the yeast. This process was repeated for 7 cycles. After 7 cycles, the mixed culture was spread onto an SDM plate with antibiotic selection, only allowing yeast colonies to grow. Yeast colonies that formed were then tested using the cell culture supernatant technique.

## Acknowledgements

We thank Dr. Chang Liu from UC Irvine for kindly providing the OrthoRep system as a gift. This work was supported by UC Riverside start-up funds granted to Y.L.

## Author contributions

C.C., S.X., and Y.L. conceived of the project, designed the experiments, analyzed the results and wrote the manuscript. S.X. and C.C. performed the experiments.

## Additional information

Competing financial interests: The authors declare that there is no conflict of interest regarding the publication of this article.

